# Development of a fluorescence based, high-throughput SARS-CoV-2 3CL^pro^ reporter assay

**DOI:** 10.1101/2020.06.24.169565

**Authors:** Heather M. Froggatt, Brook E. Heaton, Nicholas S. Heaton

**Author notes:** Address correspondence to: Nicholas S. Heaton, Assistant Professor, Department of Molecular Genetics and Microbiology (MGM), Duke University Medical Center, 213 Research Drive, 426 CARL Building, Box 3054, Durham, NC 27710.

## Abstract

In late 2019 a human coronavirus, now known as SARS-CoV-2, emerged, likely from a zoonotic reservoir. This virus causes COVID-19 disease, has infected millions of people, and has led to hundreds of thousands of deaths across the globe. While the best interventions to control and ultimately stop the pandemic are prophylactic vaccines, antiviral therapeutics are important to limit morbidity and mortality in those already infected. At this time, only one FDA approved anti-SARS-CoV-2 antiviral drug, remdesivir, is available and unfortunately, its efficacy appears to be limited. Thus, the identification of new and efficacious antivirals is of highest importance. In order to facilitate rapid drug discovery, flexible, sensitive, and high-throughput screening methods are required. With respect to drug targets, most attention is focused on either the viral RNA-dependent RNA polymerase or the main viral protease, 3CL^pro^. 3CL^pro^ is an attractive target for antiviral therapeutics as it is essential for processing newly translated viral proteins, and the viral lifecycle cannot be completed without protease activity. In this work, we present a new assay to identify inhibitors of the SARS-CoV-2 main protease, 3CL^pro^. Our reporter is based on a GFP-derived protein that only fluoresces after cleavage by 3CL^pro^. This experimentally optimized reporter assay allows for antiviral drug screening in human cell culture at biosafety level-2 (BSL2) with high-throughput compatible protocols. Using this screening approach in combination with existing drug libraries may lead to the rapid identification of novel antivirals to suppress SARS-CoV-2 replication and spread.

**IMPORTANCE:** The COVID-19 pandemic has already led to more than 400,000 deaths and innumerable changes to daily life worldwide. Along with development of a vaccine, identification of effective antivirals to treat infected patients is of the highest importance. However, rapid drug discovery requires efficient methods to identify novel compounds that can inhibit the virus. In this work, we present a method for identifying inhibitors of the SARS-CoV-2 main protease, 3CL^pro^. This reporter-based assay allows for antiviral drug screening in human cell culture at biosafety level-2 (BSL2) with high-throughput compatible sample processing and analysis. This assay may help identify novel antivirals to control the COVID-19 pandemic.

## INTRODUCTION

In December 2019, a novel human coronavirus (hCoV) was identified in the Hubei Province of China (1–3). The virus, now known as SARS-CoV-2, causes the transmissible and pathogenic disease COVID-19 (4). COVID-19 has become a global pandemic and infected over 8 million people and caused ∼500,000 deaths to date (5). Current efforts to control COVID-19 are largely focused on behavioral modifications such as social distancing and the use of masks (6). These approaches attempt to slow the spread of the virus, but meaningful control of the virus will ultimately be the result of a combination of efficacious vaccines and antiviral therapeutics (7).

Antiviral therapeutics aim to disrupt the replication cycle and reduce viral load in infected individuals. Therapeutic development efforts have led to a number of candidate antiviral compounds focused mainly on two essential viral enzymes, the RNA-dependent RNA polymerase (RdRp) and the viral proteases. Remdesivir (GS-5734), recently FDA approved as an antiviral for SARS-CoV-2, targets the polymerase to suppress hCoV replication by inducing termination of RNA polymerization (8); however, the benefits of this drug in clinical trials and early use appear limited (9). Another nucleoside analogue, β-D-N^4^-hydroxycytidine (NHC; EIDD-1931), also inhibits SARS-CoV-2 polymerase activity, likely via inducing lethal mutagenesis of the viral genome (10). In addition to the RdRp, the viral proteases, which are critical to liberate individual viral proteins from the polyprotein produced by initial genome translation, present another attractive drug target. For SARS-CoV-2, lopinavir/ritonavir, a protease inhibitor combination, is shown to interact with the main coronavirus protease, known as 3CL^pro^ or M^pro^ (11); however, early clinical trial results with these compounds have shown no significant benefits to SARS-CoV-2 patients (12). More recently, structure-based design has enabled the rapid development of new antivirals targeting the SARS-CoV-2 protease, 3CL^pro^ (13–15). At this time, these newly designed compounds are in the early stages of testing. Thus, the discovery of additional effective SARS-CoV-2 antiviral drugs remains of high importance. The identification (and subsequent improvement) of novel drugs targeting SARS-CoV-2 will require robust and high-throughput screening approaches.

Here, we report the development and validation of a fluorescent reporter optimized to detect SARS-CoV-2 3CL^pro^ activity. This assay is performed in human cell culture and does not require biosafety level 3 (BSL3) containment. Our reporter is based on FlipGFP, which only fluoresces after protease mediated activation (16). We generated and tested three reporter constructs with distinct cleavage target sequences for activation by the SARS-CoV-2 3CL^pro^. We also show that the reporter with the best signal-to-noise ratio for SARS-CoV-2 is also activatable by other coronavirus 3CL^pro^ proteins across subgroups (*beta, alpha, gamma*) and host species (human, rodent, bird). Finally, we used this reporter to test the inhibition of the SARS-CoV-2 3CL^pro^ with a known coronavirus 3CL^pro^ inhibitor, GC376 (17), and then validated the correlation between reporter inhibition and inhibition of SARS-CoV-2 viral replication. These experiments together demonstrate the utility of this approach for the identification of novel antiviral drugs that target the SAR-CoV-2 main protease, 3CL^pro^.

## RESULTS

### Generation of a fluorescent SARS-CoV-2 3CL^pro^ activity reporter

In order to develop a fluorescent reporter responsive to the SARS-CoV-2 main protease, we started with the FlipGFP protein (16). FlipGFP is used to detect protease activity by expressing the GFP beta-strands β10-11 separately from, and in a conformation incompatible with, the rest of the GFP beta-barrel, β1-9 (**Fig. 1A**). A linker containing a cleavage site holds the two GFP beta strands, 10 and 11, in an inactive, parallel conformation. When the appropriate protease is present the linker containing the cleavage site is cut. This cleavage allows GFP β11 to reorient such that GFP β10 and 11 are antiparallel and able to fit into GFP β1-9, inducing fluorescence ∼100-fold over background (16).

**Figure 1.**
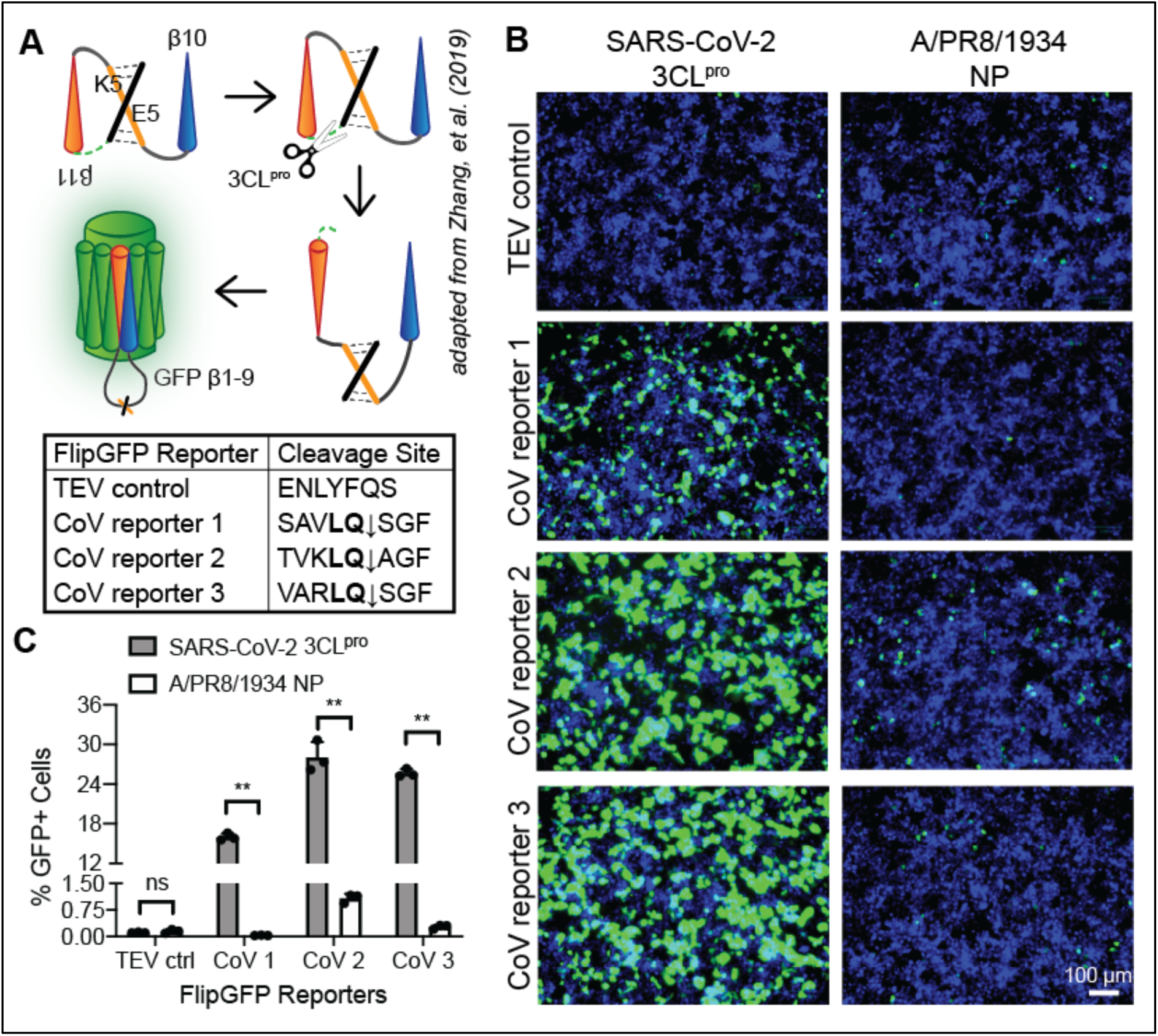
A FlipGFP protease reporter with coronavirus cleavage sites fluoresces after SARS-CoV-2 3CL^pro^ expression. A) Diagram of the FlipGFP protease reporter (16) with coronavirus cleavage sequences. FlipGFP splits GFP into β1-9 and β10-11, with β11 held in parallel to β10 by heterodimerized coiled coils E5/K5 and a linker sequence containing a coronavirus cleavage site. The CoV main protease, 3CL^pro^, cuts at the cleavage site allowing β11 to “flip” anti-parallel to β10, enabling self-assembly of the complete GFP beta-barrel and resulting in detectable fluorescence. The pan-coronavirus 3CL^pro^ consensus sequence, LQ, is in bold. Scale bars are 100μm. B) Microscopy of 293T cells 48 hours post-transfection with each FlipGFP reporter and either the SARS-CoV-2 3CL^pro^ or an influenza viral protein (A/PR8/1834 NP). Green = cleaved FlipGFP, blue = nuclei. C) Quantification of 1B. Data shown as mean ± SD, n=3, statistical analysis relative to NP control. P-values calculated using unpaired, two-tailed Student’s t-tests (*p<0.05, **p<0.001).

SARS-CoV-2 generates two proteases that cleave the viral polyprotein, a papain-like protease (PL^pro^) and a chymotrypsin-like protease (3CL^pro^). 3CL^pro^, also known as the main protease or M^pro^, is the more conserved viral protease, with only 5 amino acid changes between SARS/SARS-like CoVs and SARS-CoV-2 compared to the 102 differences found in PL^pro^ (18). 3CL^pro^ cleaves at a highly conserved consensus sequence, LQ↓, across the coronavirus family (19), and has been shown to effectively cleave luciferase-based protease biosensors (20, 21) and FRET-based assays (13, 17, 22–28). With the aim of generating a protease reporter compatible with SARS-CoV-2 and other present and future coronaviruses to support viral inhibitor screening, we selected the CoV 3CL^pro^ as our protease target.

Although the CoV 3CL^pro^ requires the minimal consensus sequence LQ↓ for cleavage, the cleavage site context influences cleavage efficiency (29). Among the CoV polyprotein cleavage sites, 3CL^pro^ targeted sites surrounding the 3CLpro sequence itself are generally cleaved most efficiently (30). Additionally, different CoVs have distinctive optimal cleavage site sequences (24). In order to develop an efficiently cleaved CoV 3CL^pro^ reporter, we tested three different cleavage sequences predicted to be highly compatible with the SARS-CoV-2 3CL^pro^ (**Fig. 1A**). CoV reporter 1 contains the conserved nsp4-5 cleavage site present in the SARS-CoV and SARS-CoV-2 viral polyproteins (31). CoV reporter 2 contains an optimized cleavage sequence for the SARS-CoV 3CL^pro^ (32). CoV reporter 3 contains an optimized sequence shown to be highly cleaved by many CoV family members (24). As a negative control, we also generated a construct harboring the Tobacco etch virus (TEV) protease cleavage site.

Our goal was to identify a construct with minimal background fluorescence while still being efficiently cleaved by SARS-CoV-2 3CL^pro^, allowing strong fluorescence for detection via microscopy, plate reader, or flow cytometry. To test our 3CL^pro^ reporters, we co-transfected each reporter with a SARS-CoV-2 3CL^pro^ expression plasmid. At 48 hours post-transfection, we could detect GFP-positive cells with each of the three CoV reporters transfected with in the SARS-CoV-2 3CL^pro^ (**Fig. 1B**). In contrast, with transfection of a negative control, nucleoprotein protein from an H1N1 influenza virus (A/PR8/1934 NP), we did not detect any signal above background levels of fluorescence. Further, the reporter containing the TEV cleavage site was not activated by SARS-CoV-2 3CL^pro^ (**Fig. 1B**). Quantification of the fluorescent signal across all of the treatment conditions demonstrated that while all three CoV reporters showed significant induction of GFP signal when co-expressed with the SARS-CoV-2 3CL^pro^, CoV reporter 2 had substantial background and the level of induction with the CoV reporter 1 reached only half of the other two CoV reporters (**Fig. 1C**). With a 100-fold change in fluorescence and minimal background, we selected CoV reporter 3 for further testing.

### Many CoV 3CL^pro^ proteins activate the FlipGFP CoV 3CL^pro^ reporter

CoV 3CL^pro^ proteins are reasonably conserved across coronavirus groups (**Fig. 2A**) (33). Further, CoV reporter 3 was based on an optimized cleavage sequence for CoV 3CL^pro^s from each coronavirus group (24). To test whether this protease reporter was compatible with a variety of CoV 3CL^pro^ proteins, we expressed CoV reporter 3 with four other coronavirus proteases from different groups (*Alphacoronavirus, Betacoronavirus* and *Gammacoronavirus*) and host species (human, mouse, bird). 48 hours after transfection, all CoV 3CL^pro^s induced visible fluorescence compared to the control influenza nucleoprotein with CoV reporter 3 (**Fig. 2B**). Quantification with a plate reader demonstrated that SARS-CoV (*Beta*, human) and avian infectious bronchitis (IBV-*Gamma*, avian) resulted in similar levels of fluorescence to SARS-CoV-2 (*Beta*, human) (**Fig. 2C**). Murine hepatitis virus (MHV-*Beta*, murine) and human coronavirus 229E (HCoV-229E-*Alpha*, human) were less compatible with CoV reporter 3, while still producing 12- and 80-fold changes in fluorescence, respectively, over background (**Fig. 2C**). These experiments show our FlipGFP 3CL^pro^ reporter is generally compatible with many CoV 3CL^pro^ proteins across coronavirus groups and host species, potentially enabling protease inhibitor screening for a variety of CoVs in addition to SARS-CoV-2.

**Figure 2.**
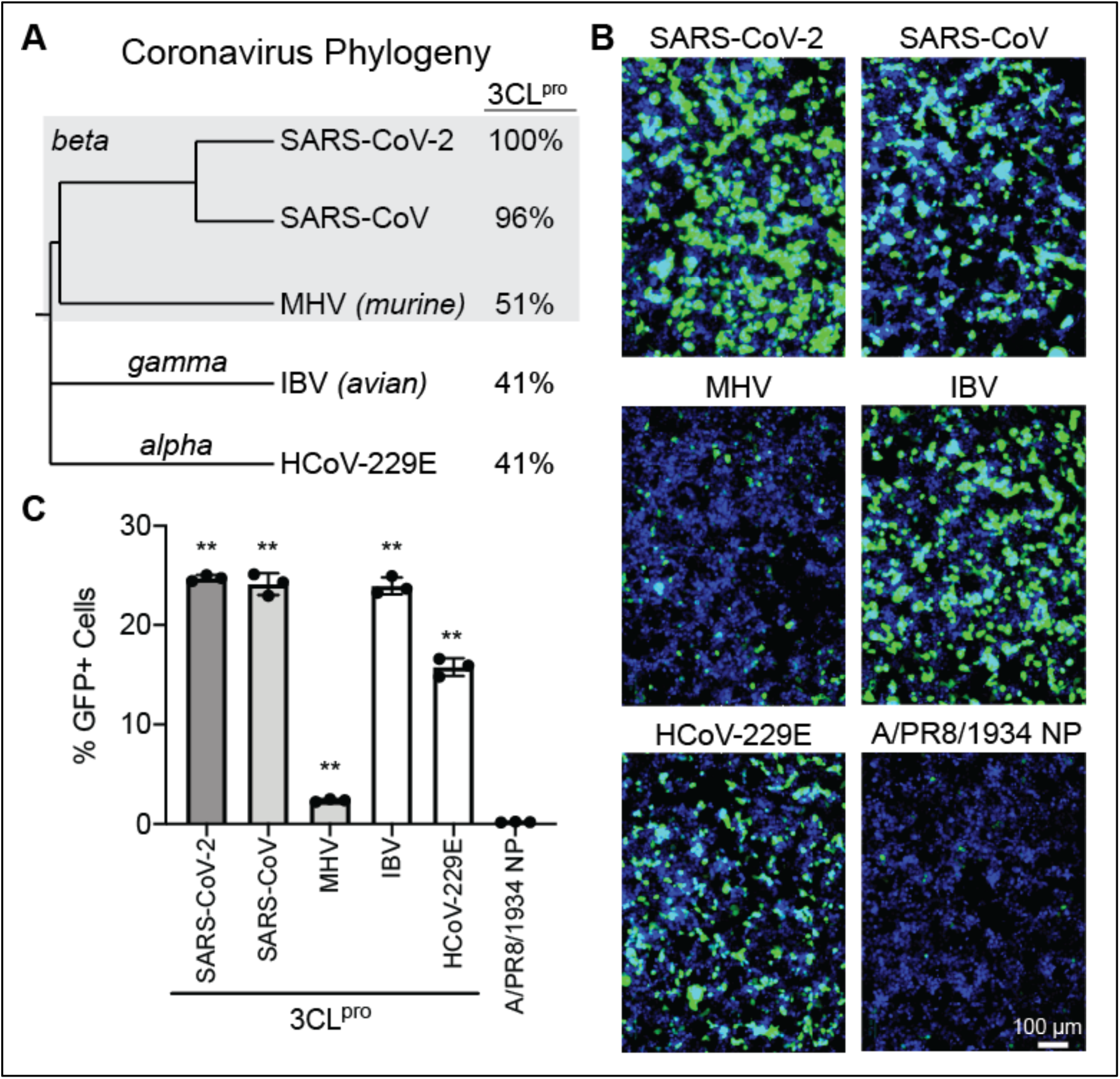
Conservation of coronavirus 3CL^pro^ activity enables CoV protease reporter compatibility with many coronaviruses. A) Phylogenetic tree of five coronaviruses, SARS-CoV-2, SARS-CoV, murine hepatitis virus (MHV), avian infectious bronchitis virus (IBV), and HCoV-229E, generated based on the polyprotein ORF1ab using NCBI Virus (34). These viruses span three coronavirus groups, *Alphacoronavirus, Betacoronavirus*, and *Gammacoronavirus*. 3CL^pro^ protein sequence identities are compared to the SARS-CoV-2 3CL^pro^. B) Microscopy of 293T cells 48 hours post-transfection with CoV reporter 3 and coronavirus 3CL^pro^ proteins or an influenza viral protein (A/PR8/1834 NP). Green = cleaved FlipGFP, blue = nuclei. Scale bars are 100μm. C) Quantification of 2B. Data shown as mean ± SD, n=3, statistical analysis relative to NP control. P-values calculated using unpaired, two-tailed Student’s t-tests (*p<0.05, **p<0.001).

### Development of a FlipGFP CoV 3CL^pro^ reporter-based assay for *in vivo* protease inhibitor screening

To develop an assay for protease inhibitor screening using our CoV reporter 3, we first needed to optimize the experimental conditions. We performed a transfection timecourse with SARS-CoV-2 3CL^pro^ to determine an early, appropriate timepoint for sample collection (**Fig. 3A**). At 12 hours post-transfection, only a few GFP fluorescing cells are visible and fluorescent signal is just above background (**Fig. 3B**). At 24 hours post-transfection, green cells are visible without appreciable background signaling (**Fig. 3B**). At 48 hours post-infection, most cells produce a high GFP signal, with some background fluorescence detectable (**Fig. 3B**). We therefore selected the 24 hour post-transfection timepoint. To increase the sensitivity of our assay, we titrated the level of SARS-CoV-2 3CL^pro^ transfected with CoV reporter 3; our goal was to maximize activation of the reporter while minimizing the amount of protease available in the cell. We transfected cells with five ratios of reporter-to-protease: 1:1, 1:0.8, 1:0.4, 1:0.2, and 1:0. 24 hours post-transfection, we observed significant decreases in reporter activation at reporter-to-protease ratios 1:0.4 and 1:0.2 (**Fig. 3C**). However, a 1:0.8 reporter-to-protease ratio resulted in no significant loss of fluorescence compared to a 1:1 ratio (**Fig. 3C**). Based on these experiments together, we selected a 1:0.8 reporter-to-protease ratio for transfection and a 24-hour post-transfection end point as the optimal conditions for our protease inhibitor assay using the FlipGFP 3CL^pro^ reporter, CoV reporter 3.

**Figure 3.**
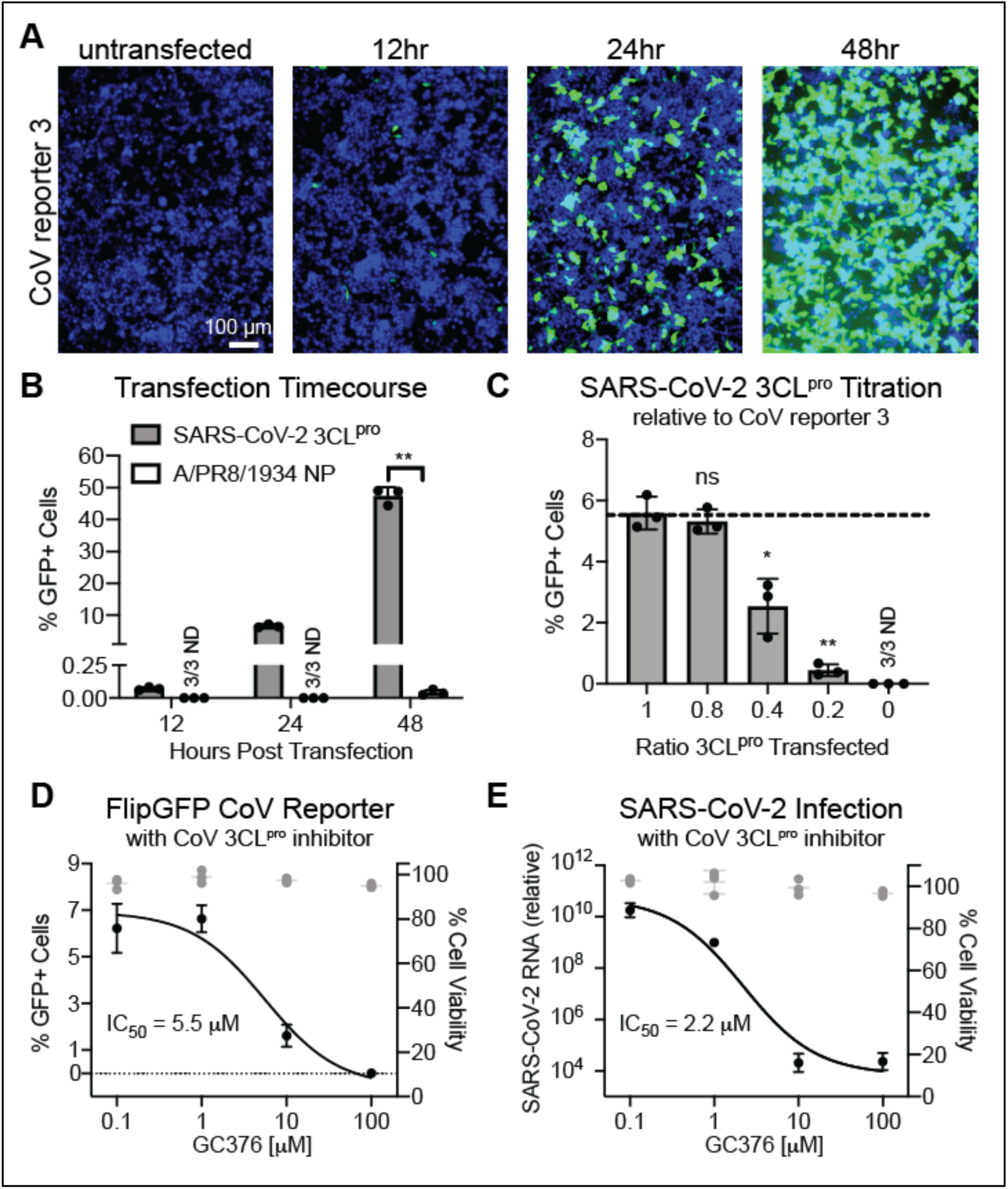
Inhibition of the SARS-CoV-2 3CL^pro^ by the protease inhibitor GC376 is measurable with the fluorescent CoV protease reporter. A) Microscopy of 293T cells before or after 12, 24, and 48 hours post-transfection with CoV reporter 3 and SARS-CoV-2 3CL^pro^. Green = cleaved FlipGFP, blue = nuclei. Scale bars are 100μm. B) Quantification of 3A. Data shown as mean ± SD, n=3, statistical analysis relative to NP control. P-values calculated using unpaired, two-tailed Student’s t-tests (*p<0.05, **p<0.001). C) Quantification of 293T cells 24 hours post-transfection with CoV reporter 3 and SARS-CoV-2 3CL^pro^, with decreasing levels of 3CL^pro^. Data shown as mean ± SD, n=3, statistical analysis relative to 1:1 ratio reporter-to-protease. P-values calculated using unpaired, two-tailed Student’s t-tests (*p<0.05, **p<0.001). D) In black: quantification of 293T cells 24 hours post-transfection with CoV reporter 3 and SARS-CoV-2 3CL^pro^ and treated with the pan-coronavirus protease inhibitor, GC376. Data shown as mean ± SD with nonlinear fit curve, n=3. In gray: cell viability was calculated relative to untransduced, vehicle-only (DMSO) samples. Data shown as mean ± SD, n=3. E) In black: RT-qPCR of VeroE6 cells 24 hours post-infection with SARS-CoV-2 (MOI 0.01) and treatment with the pan-coronavirus protease inhibitor, GC376. Data shown as mean ± SD with nonlinear fit curve, n=4. In gray: cell viability was calculated relative to un-transduced, vehicle-only (DMSO) samples. Data shown as mean ± SD, n=3.

Finally, we wanted to verify that our assay could detect drug inhibition of the SARS-CoV-2 3CL^pro^ with a known inhibitor. Therefore, we selected a recognized pan-coronavirus 3CLpro inhibitor, GC376, to test our assay (17). Four concentrations of GC376, that did not significantly impact cell viability compared to vehicle alone, were applied to cells at the time of transfection with CoV reporter 3 and SARS-CoV-2 3CL^pro^. As expected, reporter activity levels were maintained at the lower protease inhibitor concentrations, while fluorescence was reduced at the higher concentrations of GC376 (**Fig. 3D**). Thus, our assay successfully detected inhibition of SARS-CoV-3 3CL^pro^ by the protease inhibitor GC376. However, it is also important to verify that inhibition of our reporter is strongly correlated with inhibition of the SARS-CoV-2 virus. We infected VeroE6 cells with SARS-CoV-2 at an MOI of 0.01 before applying protease inhibitor at the same four concentrations as tested with the protease reporter. 24 hours post infection, we collected RNA and performed RT-qPCR to detect SARS-CoV-2 viral RNA; similar to the reporter, viral RNA levels were suppressed in a dose-dependent manner (**Fig. 3E**). Our observed inhibition of the virus is consistent with reports of inhibition of SARS-CoV-2 by GC376 in the literature (22, 23). All together, these experiments demonstrate feasibility of using our FlipGFP CoV 3CL^pro^ reporter assay to identify protease-targeting inhibitors of SARS-CoV-2.

## DISCUSSION

Our goal for this study was to develop a cell-based assay to screen for novel SARS-CoV-2 antiviral drugs at BSL2; to our knowledge, no such assay optimized for SARS-CoV-2 currently exists. Therefore, we generated a reporter requiring a coronavirus protease, 3CL^pro^, for activation of a GFP-fluorescent signal. We showed this reporter is responsive to the SARS-CoV-2 3CL^pro^, in addition to many different coronavirus 3CL^pro^ proteins. After optimizing screening conditions, we demonstrated that our reporter was sensitive to treatment with a known coronavirus protease inhibitor, GC376. These experiments illustrate the utility of our approach to identify, and subsequently optimize, novel protease inhibitors of SARS-CoV-2.

To meet the demands of virus research during the SARS-CoV-2 pandemic, reporter assays need to be flexible and high-throughout. Our reporter is activated with expression of a single CoV protein, 3CL^pro^, allowing for SARS-CoV-2 drug testing at BSL2. Additionally, the reporter is compatible with many CoV 3CL^pro^ proteins, supporting rapid testing of inhibitors against a variety of coronaviruses, present or future, and without synthesis of protease substrates or purification of viral proteins (13, 17, 22–28). Further, as our assay is performed in living cells, our system enables the discovery of protease inhibitors while simultaneously evaluating effects on cellular viability. Our assay is scalable, and the analysis requires only a basic fluorescent plate reader, supporting high-throughput screening. In addition to applications in drug discovery pipelines, this assay could be deployed to determine targets of antivirals identified via viral screening.

Reporter assays, including ours, also have limitations. Our reporter utilizes the CoV 3CL^pro^ expressed alone; during a CoV infection the protease is only one of dozens of viral proteins present, and any inhibitors that may affect cross-viral protein interactions would be missed. Additionally, CoV infection induces significant cellular membrane rearrangements that transfection of the protease alone does not. Thus, the subcellular access of therapeutic compounds to the viral protease may fail to be reflected in our assay, and the effects of an identified protease inhibitor could significantly differ when applied to authentic viral infection. Finally, although this plasmid-based expression presents many advantages, it also necessitates further screen hit testing in the context of coronavirus infection. Although the correlation between our reporter and inhibition of viral infection was appreciable with the drug GC376, testing of more inhibitors is required to make generalizable correlations between our reporter assay and viral infection readouts.

To have the greatest impact on the COVID-19 pandemic, an effective SARS-CoV-2 antiviral needs to be identified as early as possible. Countries around the world have taken drastic and necessary steps to limit the spread of virus, yet infection rates continue to rise in some. An antiviral treatment is unlikely to stop the spread of infection, but it is likely to limit the mortality associated with SARS-CoV-2 infection. It is our hope that this reporter assay facilitates the identification of SARS-CoV-2 protease inhibitor candidates to be rapidly optimized and translated to clinical use.

## MATERIALS AND METHODS

### Cell culture

All cells were obtained from ATCC and grown at 37°C in 5% CO_2_. 293T cells were grown in Dulbecco’s Modified Eagle Medium (DMEM) supplemented with 5% fetal bovine serum, GlutaMAX, and penicillin-streptomycin. VeroE6 cells were grown in MEM supplemented with 10% fetal bovine serum, pyruvate, NEAA and penicillin-streptomycin. For transfection, 24 hours before plates were poly-lysine treated and seeded with 293Ts. The next day, plasmid DNA, Opti-MEM, and TransIT-LT (Mirus) were combined using pipetting and incubated at room temperature for 20min before being added to cells by droplet.

### Plasmids

All constructs were cloned into the pLex expression vector using the BamHI and NotI restriction sites and DNA assembly (NEB). Coronavirus 3CL^pro^ expression plasmids (SARS-CoV-2, SARS-CoV, MHV, IBV, HCoV-229E) were generated using codon-optimized gBlocks (IDT). The TEV control FlipGFP was designed to include a silent NheI restriction site ahead of the TEV cleavage sequence and generated using a gBlock (IDT). The coronavirus 3CL^pro^ FlipGFP reporters (CoV reporter 1, CoV reporter 2, CoV reporter 3) were generated using forward primers containing the cleavage sequences along with the initial FlipGFP reverse primer, and assembled into the NheI and NotI digested control FlipGFP plasmid. DNA was transformed into NEB 5-alpha high efficiency competent cells (NEB). Insert size was verified with PCR and purified plasmids were sequenced using Sanger sequencing.

### Imaging and Quantification

Cells were fixed with 2% PFA at room temperature for 20 min before incubating in 1:10,000 Hoescht 33342 (Life Technologies) in PBS at 4°C overnight. Images were obtained using the ZOE Fluorescent Cell Imager (Bio-Rad). Quantification was performed using the CellInsight CX5 (Thermo Scientific).

### Cytotoxicity Assays

24 hours before treatment, plates were poly-lysine treated and seeded with 293T or VeroE6 cells. The next day, media was exchanged for complete media containing GC376 (MedKoo) at the indicated concentrations using a constant level of vehicle (DMSO). After 24 hours of treatment, cells were collected according the CellTiter-GLO (Promega) protocol and luminescence levels assessed using a luminometer.

### SARS-CoV-2 Infections and Viral qPCR

24 hours before infection, plates were poly-lysine treated and seeded with VeroE6 cells. The next day, the cells were washed with PBS before infection with SARS-CoV-2 isolate USA-WA1/2020 from BEI Resources in 2% FBS MEM infection media at an MOI of 0.01 for 1 hour. Virus was removed and cells were then placed in infection media containing GC376 (MedKoo) at the indicated concentrations using a constant level of vehicle (DMSO). 24 hours post-infection, cells were collected in TRIzol (Invitrogen) followed by RNA isolation. One-step RT-qPCR was performed with primers targeting the SARS-CoV-2 N region (BEI) using the EXPRESS One-Step Superscript qRT-PCR Kit (ThermoFisher) on an Applied Biosystems QuantStudio3 instrument. RNA was normalized using an endogenous 18S rRNA primer/probe set (Applied Biosystems).

## ACKNOWLEDGEMENTS

NSH is partially funded by the Defense Advanced Research Projects Agency’s (DARPA) PReemptive Expression of Protective Alleles and Response Elements (PREPARE) program (Cooperative agreement #HR00111920008). The views, opinions and/or findings expressed are those of the authors and should not be interpreted as representing the official views or policies of the U.S. Government. The following reagent was deposited by the Centers for Disease Control and Prevention and obtained through BEI Resources, NIAID, NIH: SARS-Related Coronavirus 2, Isolate USA-WA1/2020, NR-52281. Biocontainment work was performed in the Duke Regional Biocontainment Laboratory, which received partial support for construction from the National Institutes of Health, National Institute of Allergy and Infectious Diseases (UC6-AI058607). We would like to thank Dr. Clare Smith for help establishing SARS-CoV-2 viral infection assays at BSL3 and Laura Froggatt for designing the FlipGFP diagram.

